# Ubiquitin E3 ligases Atrogin-1 and MuRF1 protein contents are differentially regulated in the rapamycin-sensitive mTOR-S6K1 signaling pathway in C2C12 myotubes

**DOI:** 10.1101/2021.10.15.463676

**Authors:** Yusuke Nishimura, Ibrahim Musa, Peter Dawson, Lars Holm, Yu-Chiang Lai

## Abstract

Muscle-specific ubiquitin E3 ligases, Atrogin-1 and MuRF1, are highly expressed in multiple conditions of skeletal muscle atrophy. The PI3K/Akt/FoxO signaling pathway is well known to regulate Atrogin-1 and MuRF1 gene expressions. Evidence supporting this is largely based on stimuli by insulin and IGF-1, that activate anabolic signaling, including Akt and Akt-dependent transcription factors. However, Akt activation also activates the mammalian target of rapamycin complex 1 (mTORC1) which induces skeletal muscle hypertrophy. However, whether mTORC1-dependent signaling has a role in regulating Atrogin-1 and/or MuRF1 gene and protein expression is currently unclear. In this study, we confirmed that activation of insulin-mediated Akt signaling suppresses both Atrogin-1 and MuRF1 protein content and that inhibition of Akt increases both Atrogin-1 and MuRF1 protein content in C2C12 myotubes. Interestingly, inhibition of mTORC1 using a specific mTORC1 inhibitor, rapamycin, increased Atrogin-1, but not MuRF1, protein content. Furthermore, activation of AMP-activated protein kinase (AMPK), a negative regulator of the mTORC1 signaling pathway, also showed distinct time-dependent changes between Atrogin-1 and MuRF1 protein content, suggesting differential regulatory mechanisms between Atrogin-1 and MuRF1 protein content. To further explore the downstream of mTORC1 signaling, we employed a specific S6K1 inhibitor, PF-4708671, and found that Atrogin-1 protein content was dose-dependently increased with PF-4708671 treatment, whereas MuRF1 protein content was not significantly altered. Overall, our results indicate that Atrogin-1 and MuRF1 protein contents are regulated by different mechanisms, the downstream of Akt, and that Atrogin-1 protein content can be regulated by rapamycin-sensitive mTOR-S6K1 dependent signaling pathway.

## 1. Introduction

Atrogin-1 (also known as Muscle atrophy F-box protein: MAFbx or *FBXO32*) and Muscle-specific RING finger protein 1 (MuRF1 or *TRIM63*) are muscle specific E3 ligases and their expression is highly associated with various skeletal muscle atrophic models [1, 2]. In agreement with above, a plethora of studies have confirmed that Atrogin-1 and MuRF1 mRNA expression are useful molecular biomarkers of skeletal muscle atrophy [3]. Although both Atrogin-1 and MuRF1 gene expressions increase in almost all atrophic models, various muscle atrophic conditions (e.g., fasting, immobilization, diabetes, insulin resistance) are likely to alter multiple signaling pathways to control Atrogin-1 and MuRF1 gene and protein expression [4]. While the PI3K-Akt signaling pathway is known to regulate Atrogin-1 and MuRF1 gene expression, the mechanisms that regulate protein content of these two E3 ligases remain to be elucidated. Many studies have assumed that mRNA expressions implicitly reflect the corresponding changes of protein content, but in reality the expression levels of individual mRNA and its corresponding protein are indeed poorly correlated [5, 6]. The poor correlation can be explained by multiple processes, including transcription and degradation of mRNAs, translation, folding, and degradation of proteins [7, 8]. As protein is the final product executing gene function, direct measurement of protein content should be more relevant to biological functions [8, 9]. However, in the cases of Atrogin-1 and MuRF1, poor quality of antibodies is often a major obstacle to reveal protein content in biological samples [3, 10, 11].

PI3K/Akt/forkhead box (FoxO) signaling is one of the most well studied pathways known to regulate Atrogin-1 and MuRF1 mRNA transcription expression [12-14]. Studies have shown that treatment of IGF-1 or an introduction of constitutively active Akt prevents both Atrogin- 1 and MuRF1 mRNA transcription expression in C2C12 myotubes [12, 13]. In addition, denervation-induced skeletal muscle atrophy was prevented by IGF-1 treatment [13]. IGF-1 increases Akt phosphorylation and suppresses Atrogin-1 and MuRF1 mRNA transcription expression in mouse skeletal muscle [13], indicating a link between Akt and Atrogin-1/MuRF1 axis in skeletal muscle atrophy. Mechanistically, Akt phosphorylates the transcription factor FoxO to induce FoxO nuclear exclusion, which downregulates FoxO-dependent gene transcription [15]. A study has also confirmed that overexpression of FoxO3a in mouse skeletal muscle is able to induce Atrogin-1 mRNA expression and an atrophic phenotype [12]. In contrast, siRNA knockdown of FoxO1-3 inhibits Atrogin-1 promoter activity measured by Atrogin-1 luciferase reporter constructs during fasting-induced muscle atrophy [12]. All these findings have evidenced that Akt-FoxO axis is critical for regulating Atrogin-1 and MuRF1 mRNA transcriptional expression. However, some contradictory results have also been reported. For example, a study showed that deletion of Akt1 or Akt2 did not alter Atrogin-1 mRNA and protein expressions in mouse skeletal muscle [16]. Atrogin-1 and MuRF1 mRNA expression, including Atrogin-1 protein content, were shown to be unchanged in ageing-induced muscle atrophy, where Akt activity and FoxO3a phosphorylation were elevated, compared to young control skeletal muscles [17]. These contradictory findings raise the question of whether Akt-FoxO axis is the solely signaling pathway that regulate Atrogin-1 and MuRF1 expression.

mTORC1 plays an important role in regulating protein synthesis and autophagy-lysosome system [18], and its activation has been well associated with skeletal muscle hypertrophy [19, 20]. Surprisingly, the involvement of mTORC1 in regulating muscle protein degradation has not been well investigated. A recent study led by Zhao et al. [21] suggested that mTOR (including mTORC1 and mTORC2) may be involved in the regulation of protein degradation in C2C12 myotubes. Their previous study has shown that treatment of rapamycin, a specific mTORC1 inhibitor, can increase protein degradation in C2C12 myotubes [22], which lead the authors to suggest mTORC1 as a contributor for controlling protein degradation. Furthermore, there is also evidence suggesting that Atrogin-1 and MuRF1 mRNA expressions are regulated by distinct signaling mechanisms. Sacheck et al. [22] showed that rapamycin treatment increases Atrogin-1, but not MuRF1, mRNA expression. However, proof at protein level is currently lacking and such information is needed to better understand what the signaling mechanisms are controlling Atrogin-1 and MuRF1 protein content, which essentially execute the enzymatic ubiquitin E3 ligase activity.

The present study therefore aims to investigate whether the downstream of Akt, such as mTORC1 and S6K1 signaling pathway, is involved in controlling Atrogin-1 and MuRF1 protein content in C2C12 myotubes. Using small molecules inhibiting mTORC1 or S6K1 activity, we demonstrated that Atrogin-1, but not MuRF1, protein content is regulated in the rapamycin-sensitive mTOR and S6K1-dependent signaling pathways. Our results suggest that the role of Akt-FoxO is not the only signaling pathway regulating Atrogin-1 protein content and that the downstream of Akt, such as the rapamycin-sensitive mTOR and S6K1-dependent signaling pathways, are involved in regulating Atrogin-1 protein content in skeletal muscle.

## 2. MATERIALS AND METHODS

### 2.1 C2C12 cell culture

Mouse skeletal muscle C2C12 myoblast cells were obtained from the American Type Culture Collection (ATCC, Manassas, VA, USA). Cells were seeded and maintained in Dulbecco’s Modified Eagle Medium (DMEM; Thermo Fisher Scientific, Loughborough, UK, 31966021) containing GlutaMAX, 25 mM of glucose, and 1 mM of sodium pyruvate, supplemented with 10% (v/v) of Hyclone fetal bovine serum (FBS, Fisher Scientific, Loughborough, UK, SV30180.03), 1% (v/v) of Penicillin-Streptomycin (10 000 Units/mL-ug/mL, Thermo Fisher Scientific, Loughborough, UK, 15140122). Myoblasts were seeded onto six-well multidishes (greiner bio-one, 657 160) and when confluency reached at 90%, myoblasts were differentiated into myotubes in DMEM supplemented with 2% (v/v) of horse serum (Sigma-Aldrich, Cambridgeshire, UK, H1270), 1% (v/v) of Penicillin-Streptomycin. The media was changed every 48 h. Cultures were maintained in a humified incubator at 37 □ with an atmosphere of 5% of CO2 and 95% of air.

### 2.2 Drug reconstitution and cell treatment

Akt1/2/3 inhibitor MK-2206 dihydrochloride (ApexBio, A3010), Rapamycin (Sigma-Aldrich, 553211), adenosine monophosphate (AMP)–activated protein kinase (AMPK) activator 991 (AOBIOUS, MA, USA, AOB8150), S6K1 Inhibitor, PF-4708671 (Sigma-Aldrich, Dorset, UK, 559278) were prepared in DMSO and treatment conditions were described in the figure legend. Insulin solution human was obtained from Sigma (Sigma Aldrich, Dorset, UK, I9278).

### 2.3 Cell lysis

Cells were lysed in ice-cold sucrose lysis buffer containing: 250 mM of sucrose, 50 mM of Tris-base (pH 7.5), 50 mM of sodium fluoride, 10 mM of sodium β-glycerophosphate, 5 mM of sodium pyrophosphate, 1 mM of EDTA, 1 mM of EGTA, 1 mM of benzamidine, 1 mM of sodium orthovanadate, 1 x complete Mini EDTA-free protease inhibitor cocktail (Roche), 1% of Triton X-100, and 100 mM of 2-chloroacetamide. Cell lysates were centrifuged for 15 minutes at 13 000 g at 4°C and the supernatant was stored at -80°C before analysis for total protein concentrations using the Bradford protein assay (Thermo Fisher Scientific, Leicestershire, UK, 23200). Protein in each sample was quantified from a standard curve using BSA standards (Thermo Fisher Scientific, Leicestershire, UK, 23209).

### 2.4 Western blot

Cell lysates were prepared in 1x NuPAGE LDS sample buffer (Invitrogen, NP0008) containing 2-mercaptoethanol (final concentration 1.5%) and left to denature overnight at room temperature. Prepared cell lysates (10-15 μg of total protein) were loaded into 8% or 10% Bis/Tris gels prior to sodium dodecyl sulfate– polyacrylamide gel electrophoresis (SDS-PAGE). Gels were run in 1x MOPS buffer for approximately 60 minutes at 140V. Proteins were transferred onto 0.2 μm polyvinylidene fluoride (PVDF) membranes (Millipore, Hertfordshire, UK) for 1 hour at 100V. Membranes were blocked in 5% of milk diluted in Tris-buffered saline Tween-20 (TBS-T): 137 mM of sodium chloride, 20 mM of Tris-base 7.5 pH, 0.1% of Tween-20 for 1 hour. After blocking, membranes were washed 3 times for 5 min in TBS-T before being incubated overnight at 4°C with the appropriate primary antibodies (**Table 1**). Membranes were washed 3 times for 5 min in TBS-T prior to incubation in horse radish peroxidase-conjugated secondary antibodies (see Supplementary Table 1) at room temperature for 1 h. Membranes were washed a further three times in TBS-T prior to antibody detection using enhanced chemiluminescence horseradish peroxidase substrate detection kit (Millipore, Hertfordshire, UK). Imaging was undertaken using a G:BOX Chemi-XR5 (Syngene, Cambridgeshire, UK). Band intensities were quantified using ImageJ/Fiji (NIH, Bethesda, MD, USA). Phosphorylation levels were determined by the expression of phosphorylated protein divided by expression of non-phosphorylated total protein. Vinculin was used as the loading control.

**Table 1.**
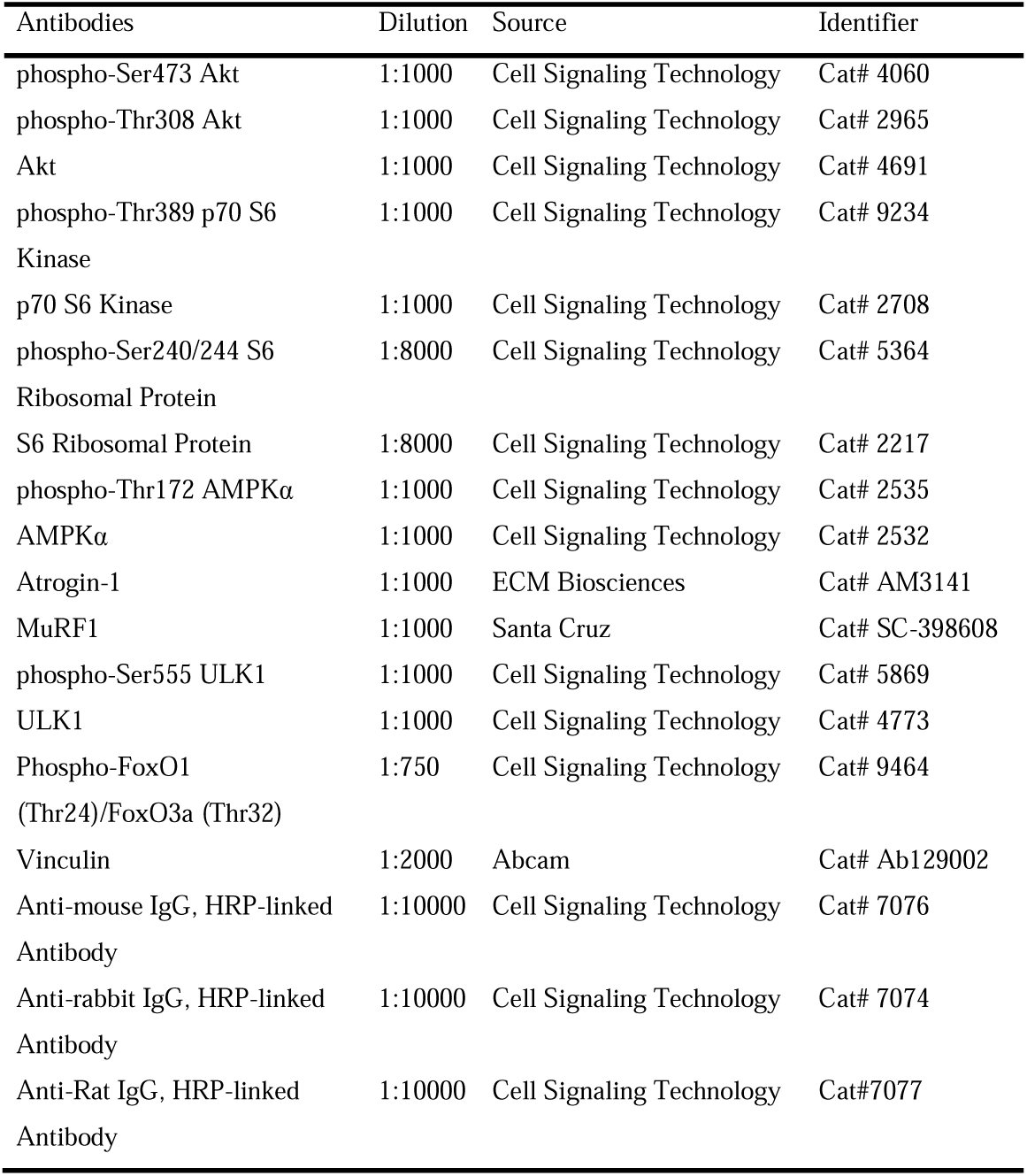
Antibodies for western blot.

### 2.5 Statistical analysis

The statistical analyses were performed using Prism version 8.1.2 (GraphPad Software, San Diego, California USA, www.graphpad.com). Values of *P* < 0.05 (*) were considered statistically significant. For time course and dose-response experiments, a one-way analysis of variance (ANOVA) was performed with Dunnett’s post-hoc test compared to control (CON). Data are presented as mean ± SD. All experiments were performed in duplication and repeated at least twice.

## 3. Results

### 3.1 Evidence of Insulin/Akt/FoxO signaling pathway modulating Atrogin-1 and MuRF1 protein content

We first confirmed if insulin/Akt/FoxO signaling pathway is sufficient to modulate both Atrogin-1 and MuRF1 protein content in C2C12 myotubes. Using an allosteric Akt inhibitor (MK-2206), we showed that Atrogin-1 protein content was significantly increased at 3 h, 6 h, and 9 h after the treatment of 10 μM MK-2206 (**Fig. 1** B). MuRF1 protein content was also significantly increased at 6 h after the treatment of MK-2206 (**Fig. 1** C). In line with a previous study [23], Akt phosphorylation at Ser^473^ and Thr^308^ was completely abolished over the course of 9 h treatment with MK-2206 (**Fig. 1** A). We also confirmed that inhibition of Akt activity prevents FoxO1 and FoxO3a phosphorylation and reduces S6K1 and rpS6 phosphorylation (**Fig. 1** A).

**Figure 1.**
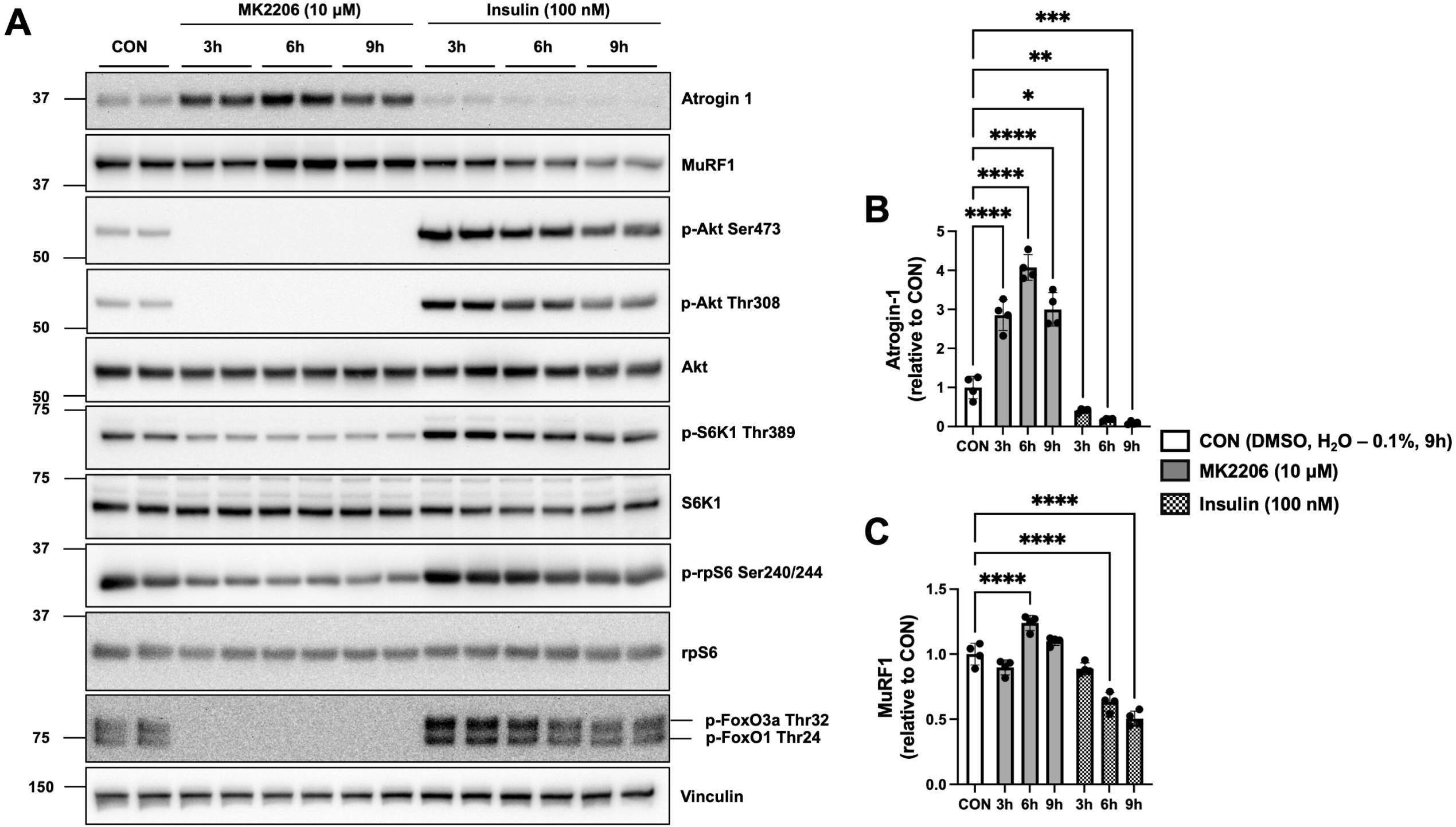
Insulin/Akt signaling pathway is sufficient to modulate Atrogin-1 and MuRF1 protein contents in C2C12 myotubes. C2C12 myotubes were treated with DMSO (0.1%, 9 h) as a vehicle control (CON), MK2206 (10 μM), or insulin (100 nM) for 3, 6, or 9 h. Lysates were analyzed by SDS-PAGE and western blotting with the indicated antibodies. (A) Representative images from one of two independent experiments. (B) Quantification of Atrogin-1. (C) Quantification of MuRF1. Data are expressed as means ± SD (n = 4) fold changes relative to CON. One-way ANOVA with Dunnett’s post-hoc test, \**P* < 0.05, \*\**P* < 0.01, \*\*\**P* < 0.001, \*\*\*\**P* < 0.0001 compared to CON.

Atrogin-1 protein content was significantly decreased at 3 h, 6 h, and 9 h following the treatment of 100 nM insulin stimulation (**Fig. 1** B). MuRF1 protein content was also significantly decreased at 6 h and 9 h after insulin treatment (**Fig. 1** C). As expected, insulin stimulated Akt phosphorylation at both Ser^473^ and Thr^308^ sites. The enhanced Akt activity was also confirmed by the increases of its downstream, such as FoxO1, FoxO3a, S6K1, and rpS6 phosphorylation (**Fig. 1** A).

### 3.2 Atrogin-1, but not MuRF1, protein content is increased by the rapamycin sensitive mTORC1 inhibition

Treatment with Rapamycin can specifically inhibit mTORC1 activity without directly affecting mTORC2 activity, but a long-term treatment (≥ 24 h) is known to inhibit mTORC2 activity [24]. Therefore, we have limited the treatment time of small molecules to not more than 9 h. Interestingly, Atrogin-1 protein content was increased at 3 h, 6 h, and 9 h following the treatment of 100 nM rapamycin (**Fig. 2** B). Despite that Atrogin-1 protein content was increased, MuRF1 protein content remained unchanged (**Fig. 2** C). As anticipated, rapamycin treatment completely inhibited S6K1 and rpS6 phosphorylation (**Fig. 2** A) without inducing a significant change in Akt phosphorylation (*P* = 0.38).

**Figure 2.**
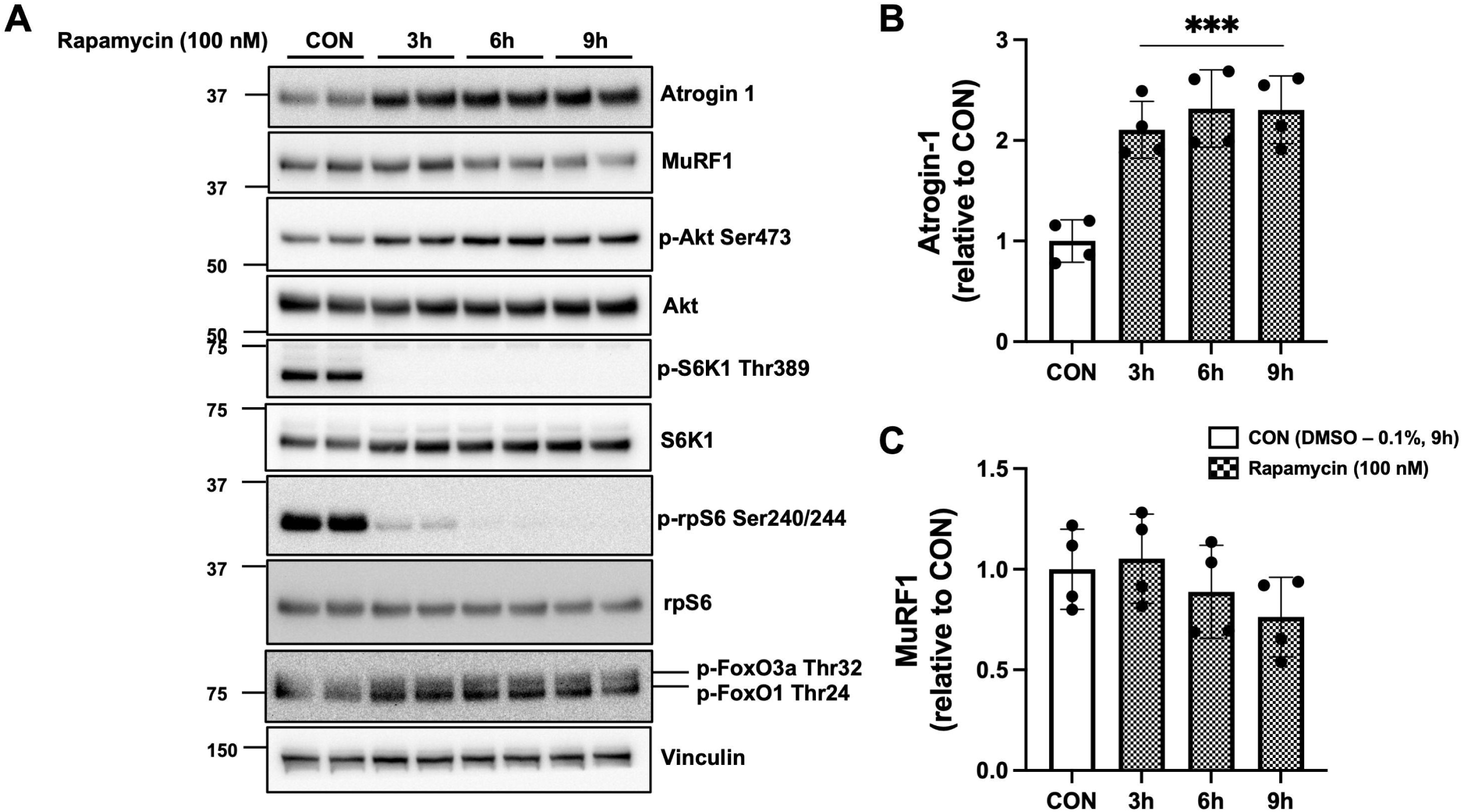
Rapamycin-sensitive mTOR inhibition increases Atrogin-1, but not MuRF1 protein contents in C2C12 myotubes. C2C12 myotubes were treated with DMSO (0.1%, 9 h) as a vehicle control (CON) or Rapamycin (100 nM) for 3, 6, or 9 h. Lysates were analyzed by SDS-PAGE and western blotting with the indicated antibodies. (A) Representative images from one of two independent experiments. (B) Quantification of Atrogin-1. (C) Quantification of MuRF1. Data are expressed as means ± SD (n = 4) fold changes relative to CON. One-way ANOVA with Dunnett’s post-hoc test, \*\*\**P* < 0.001 compared to CON.

### 3.3 Distinct time-dependent changes of Atrogin-1 and MuRF1 protein content following AMPK activation

AMPK activation is known to inhibit mTORC1 activity [25] via the phosphorylation of tuberous sclerosis complex 2 (TSC2) [26] and Raptor [27]. To further investigate the role of mTORC1 on the regulation of Atrogin-1 and MuRF1 protein content, we used a direct AMPK activator, 991, to increase AMPK activity in C2C12 myotubes [25, 28]. Interestingly, Atrogin-1 protein content was increased rapidly at 3 h and 6 h, despite returning to the basal level after 9 h of treatment (**Fig. 3** B). In contrast, MuRF1 protein content had obviously delayed increases at 6 h and 9 h after 991 treatment (**Fig. 3** C). These results again suggest that Atrogin-1 and MuRF1 protein content are regulated by distinct signaling mechanisms. As expected, ULK1 phosphorylation at Ser^555^ was increased by the treatment of 991 (**Fig. 3** A) [29] and the inhibition of mTORC1 activity was confirmed by showing a decrease in S6K1 and rpS6 phosphorylation (**Fig. 3** A).

**Figure 3.**
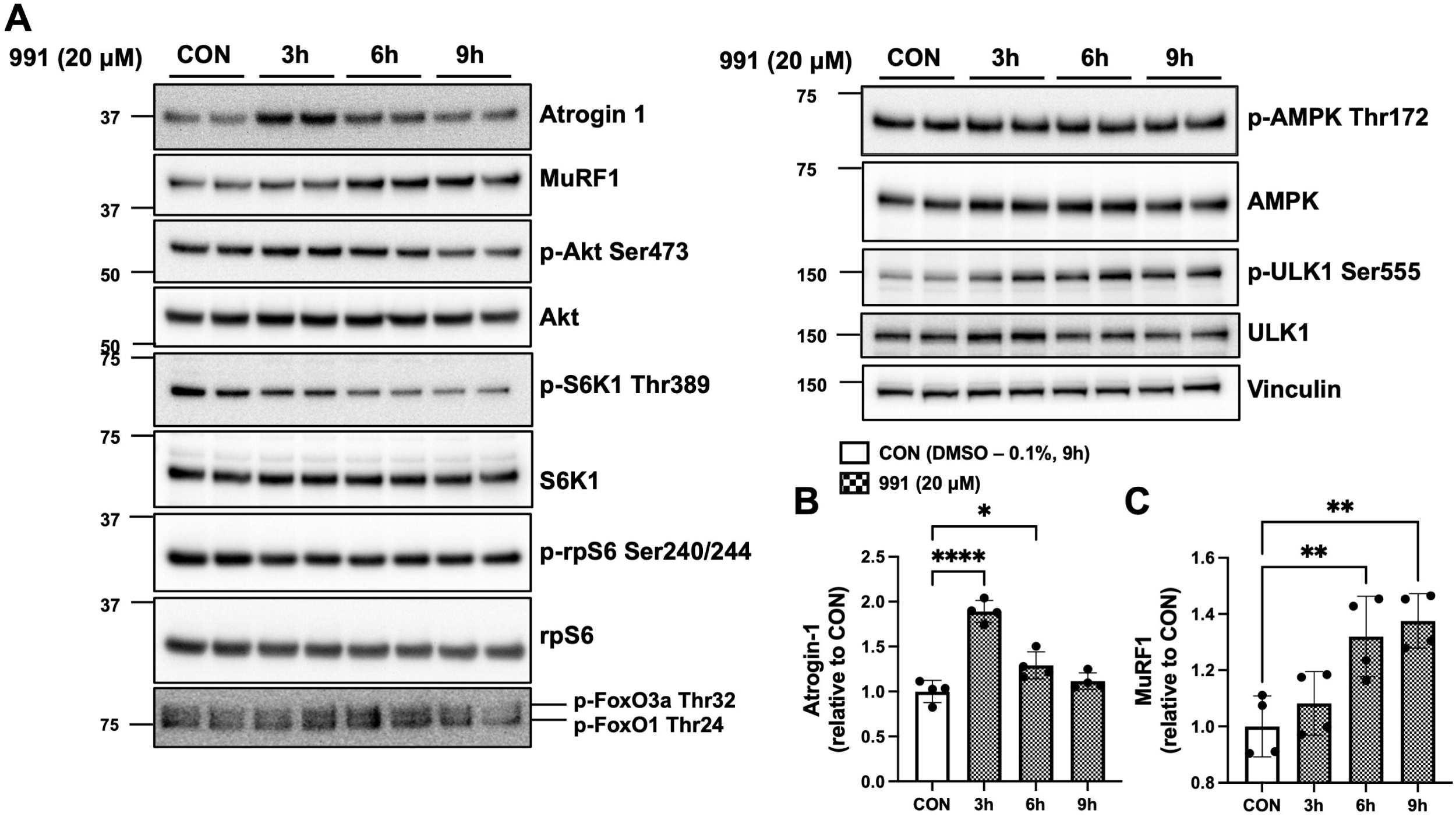
Inhibition of the mTORC1 pathway by AMPK activator 991 on Atrogin-1 and MuRF1 protein contents in C2C12 myotubes. C2C12 myotubes were treated with DMSO (0.1%, 9 h) as a vehicle control (CON) or 991 (20 μM) for 3, 6, or 9 h. Lysates were analyzed by SDS-PAGE and western blotting with the indicated antibodies. (A) Representative images from one of two independent experiments. (B) Quantification of Atrogin-1. (C) Quantification of MuRF1. Data are expressed as means ± SD (n = 4) fold changes relative to CON. One-way ANOVA with Dunnett’s post-hoc test, \**P* < 0.05, \*\**P* < 0.01, \*\*\*\**P* < 0.0001 compared to CON.

### 3.4 Atrogin-1 protein content is increased by S6K1 inhibition

To further explore the distinct mechanisms that regulate Atrogin-1 and MuRF1 protein content, we asked whether mTORC1 downstream, such as S6K1, is involved in regulating Atrogin-1 or MuRF1 protein content. Using a specific S6K1 inhibitor [30], we showed that Atrogin-1 (**Fig. 4** B) protein content was increased in a dose-response manner, where significant increases was seen with the treatment of 40 μM and 50 μM PF-4708671. Instead of increasing, MuRF1 protein content was indeed decreased at 50 μM (**Fig. 4** C). Inhibition of S6K1 was confirmed by the observation of reduced rpS6 phosphorylation (**Fig. 4** A). As expected, the phosphorylation of S6K1 was increased by the treatment of PF-4708671 [30] (**Fig. 4** A). Next, we performed Pearson’s correlation coefficient to identify the relationship between p-rpS6^Ser240/244^/rpS6 and Atrogin-1 or MuRF1 by plotting the dose-response data (**Fig. 4** D). Interestingly, a strong negative correlation was observed between p-rpS6^Ser240/244^/rpS6 and Atrogin-1 (r = - 0.90, *P* < 0.0001), whereas no significant association was observed between p-rpS6^Ser240/244^/rpS6 and MuRF1 (r = 0.17, *P* = 0.44).

**Figure 4.**
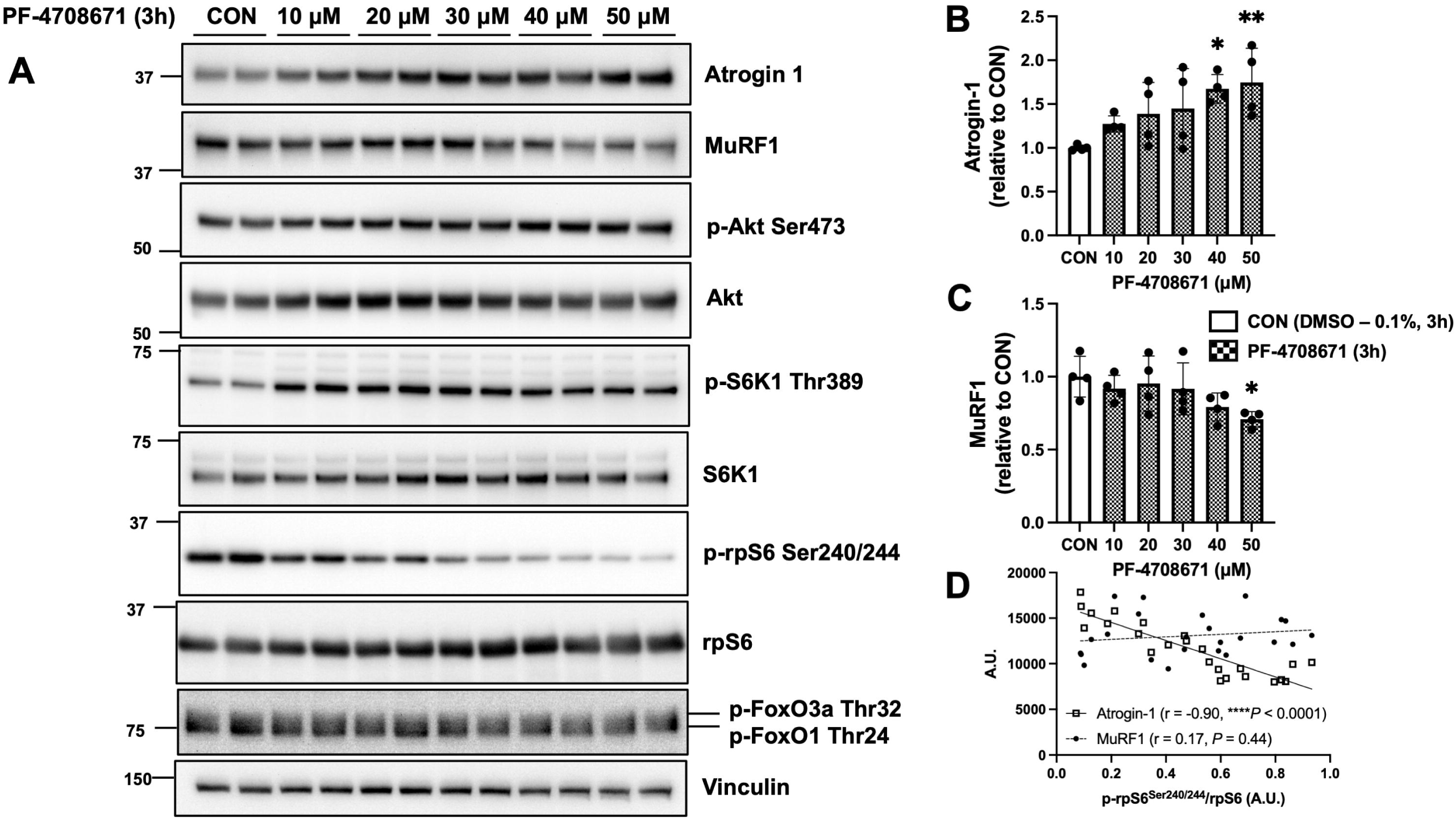
A dose-response effect of S6K1 inhibitor on Atrogin-1 and MuRF1 protein contents in C2C12 myotubes. C2C12 myotubes were treated with DMSO (0.1%, 3 h) as a vehicle control (CON) or PF-4708671 at the indicated doses for 3 h. Lysates were analyzed by SDS-PAGE and western blotting with the indicated antibodies. (A) Representative images of 2 independent experiments. (B) Quantification of Atrogin-1. (C) Quantification of MuRF1. Data are expressed as means ± SD (n = 4) fold changes relative to CON. One-way ANOVA with Dunnett’s post-hoc test, \**P* < 0.05, \*\**P* < 0.01, \*\*\**P* < 0.001, \*\*\*\**P* < 0.0001 compared to CON. (D) Pearson’s correlation coefficient to identify the association between p-rpS6^Ser240/244^/rpS6 and Atrogin-1 or MuRF1.

To confirm that S6K1 inhibition increases Atrogin-1, but not MuRF1, protein content, we performed a time course experiment using 30 μM of PF-4708671 for up to 24 h (**Fig. 5** A). As anticipated, the protein content of Atrogin-1 was increased over the course of PF-4708671 treatment at 3 h, 6 h, and 24 h (**Fig. 5** B). Although MuRF1 protein content (**Fig. 5** C) remained unchanged over majority of the time points, there was still an unexpected increase occurred at 6 h after PF-4708671 treatment.

**Figure 5.**
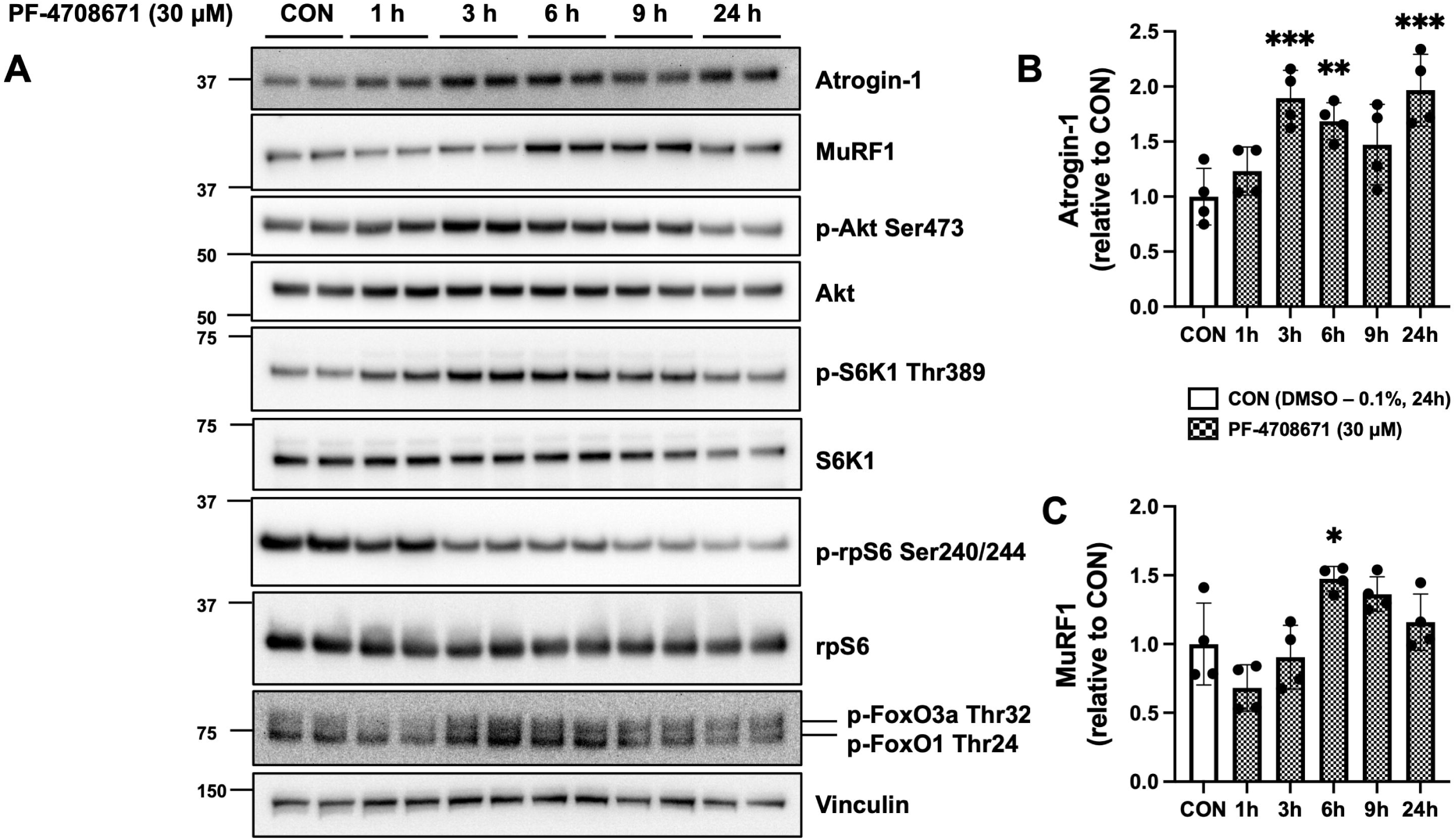
A time course effect of S6K1 inhibitor on Atrogin-1 and MuRF1 protein contents in C2C12 myotubes. C2C12 myotubes were treated with DMSO (0.1%, 24 h) as a vehicle control (CON) or PF-4708671 (30 μM) for up to 24 h. Lysates were analyzed by SDS-PAGE and western blotting with the indicated antibodies. (A) Representative images from one of two experiments. (B) Quantification of Atrogin-1. (C) Quantification of MuRF1. Data are expressed as means ± SD (n = 4) fold changes relative to CON. One-way ANOVA with Dunnett’s post-hoc test, \**P* < 0.05, \*\**P* < 0.01, \*\*\**P* < 0.001 compared to CON.

## 4. Discussion

The gene expression of Atrogin-1 and MuRF1 are highly associated with almost all kinds of skeletal muscle atrophy [1-3]. Genetic studies have also shown that knockout of Atrogin-1 or MuRF1 partially rescue denervation-induced skeletal muscle atrophy [1]. However, the molecular mechanisms of how Atrogin-1 and MuRF1 contribute to skeletal muscle atrophy are still unclear. The most recent study has indicated that the enzymatic activity of these ubiquitin E3 ligases is particularly important in controlling skeletal muscle mass [31]. Therefore, obtaining information relevant to the regulation of Atrogin-1 and MuRF1 protein content will provide an alternative opportunity to manipulate their functional E3 ligase activity. This information will also help identify new therapeutic targets to treat and/or prevent skeletal muscle atrophy. Here, we have made use of small molecules to evaluate some key signaling pathways that modulate Atrogin-1 and MuRF1 protein contents in C2C12 myotubes. In accordance with previous studies, we confirmed that insulin/Akt/FoxO pathway is sufficient to modulate both Atrogin-1 and MuRF1 protein contents, which is in agreement with the tendency of measuring mRNA transcriptional expression [12-14, 22]. Further investigation revealed that Atrogin-1, but not MuRF1, protein content is predominantly increased when rapamycin-sensitive signaling pathways is inhibited. These findings show that Atrogin-1 and MuRF1 protein contents are regulated through different mechanisms downstream of Akt. More interestingly, our studies also revealed that Atrogin-1 protein content can be regulated by S6K1 dependent signaling pathway.

Inactivation of PI3K/Akt/FoxO signaling pathway is well known as an “atrophic signal” that increases both MuRF1 and Atrogin-1 mRNA expression [32]. However, few studies have investigated whether MuRF1 and Atrogin-1 protein contents are regulated in accordance with their gene/mRNA expressions. The current study confirmed that protein content of both Atrogin-1 and MuRF1 were suppressed by insulin, whereas Atrogin-1 and MuRF1 protein contents were upregulated by the treatment of MK-2206. These findings are consistent with the mRNA expressions investigated by previous studies [12-14, 22].

In the present study, we showed that Atrogin-1 protein content was increased after 3 h treatment of rapamycin, whereas MuRF1 protein content was not changed throughout the time course (**Fig. 2**). This data indicates that inhibition of mTORC1 signaling can enhance Atrogin-1, but not MuRF1, protein content. This is indeed consistent with a previous study reported that inhibition of rapamycin-sensitive signaling pathway increases Atrogin-1, but not MuRF1, mRNA expression [22]. Our findings strengthened the previous evidence of mRNA data [22] by showing that inhibition of rapamycin-sensitive mTOR-S6K1 signaling pathway also induces an increase in Atrogin-1 protein content.

The most interesting findings in the present study are that Atrogin-1 and MuRF1 protein contents can be regulated differently, and that Atrogin-1 protein content is regulated by rapamycin-sensitive and S6K1 dependent signaling pathways. In the present study, the phosphorylation of FoxO3a at Thr^32^ and FoxO1 at Thr^24^ was not altered after rapamycin or PF-4708671 treatment, suggesting that FoxOs are not the most critical factor regulating Atrogin-1 (as well as MuRF1) protein content. Multiple transcription factors, including the NF-κB transcription factors CCAAT/enhancer-binding protein-β (C/EBPβ) and Smad3, can work cooperatively to regulate Atrogin-1 mRNA transcription Atrogin-1 [3, 33]. Thus, complex cooperative mechanisms of transcription factors might have been involved in the distinct protein expression patterns between Atrogin-1 and MuRF1 protein content. In supporting our finding that Atrogin-1 protein content is regulated by S6K1 dependent signaling, previous studies have also shown that the absence of S6K1 causes skeletal muscle atrophy in mice [34]. In addition, Marabita et al. [35] reported that S6K1 is required for the prevention of protein aggregation during skeletal muscle hypertrophy in mice. These observations led us to hypothesize that protein quality control, mainly protein degradation, is the mechanism inducing the increased Atrogin-1 protein content, when rapamycin-sensitive mTOR-S6K1 signaling is inhibited. However, future studies should confirm this hypothesis by investigating the process of Atrogin-1 protein turnover rate, and subsequent protein content.

The mTORC1 signaling pathway has been shown as a positive regulator of skeletal muscle mass in several models of hypertrophy [19, 20, 36]. In support of age-related muscle loss, studies have demonstrated that muscle contraction-induced activation of mTORC1 signaling is impaired with ageing [37, 38]. In contrast, constant activation of mTORC1 is known to cause myopathy, but not hypertrophy [39]. Moreover, a most recent study led by Joseph et al. [40] showed that mTORC1 signaling pathway is indeed hyperactivated in age-related muscle loss with a concomitant increase in both Atrogin-1 and MuRF1 mRNA expression in basal rat skeletal muscle. More interestingly, a partial inhibition of mTORC1 via RAD001 restored age-related skeletal muscle loss [40]. RAD001 treatment also decreased MuRF1 mRNA expression while Atrogin-1 mRNA was not altered in ageing muscle. We cannot directly compare our findings to their results as they did not report Akt activity and information of MuRF1 and Atrogin-1 protein contents was not available. Nonetheless, these findings indicate the importance of fine tuning the mTORC1 activity in maintaining skeletal muscle mass and Atrogin-1 and/or MuRF1 may be responsible for this.

Protein content is determined by protein turnover, which is a continuous process of protein synthesis and protein degradation [41]. In this study, Atrogin-1 and MuRF1 protein contents were investigated following time-course and/or dose-dependent small molecule treatments, which is a snapshot in time of the impact of the protein turnover kinetics on protein balance. mTORC1 is a well-known signaling pathway to control protein synthesis. Thus, after the treatment of rapamycin or PF-4708671, a decrease in protein synthesis would be expected and a greater decrease in protein degradation would, in theory, contribute to the observed increase in Atrogin-1 protein content. Post-translational modifications and the subsequent degradation make it more complicated to understand how protein content is regulated. For example, many ubiquitin E3 ligases have been implicated to regulate their own protein abundance [42] because most of E3 ligases have the ability to ubiquitylate itself (known as autoubiquitylation) and trigger self-degradation processes (either via proteasome or autophagic lysosome). For example, the greater autoubiquitylation usually demonstrates greater E3 ligase activity [11], which was observed in MuRF1 via in vitro reaction [1]. However, the degree of autoubiquitylation on MuRF1 and Atrogin-1 is currently not clear in any of muscle atrophy conditions. Although it is not clear from the present study, autoubiquitylation might have been involved in the regulation of Atrogin-1 and MuRF1 protein contents.

While our findings suggest that Atrogin-1 and MuRF1 protein contents are regulated by different signaling mechanisms, future studies should determine which molecules in the rapamycin-sensitive mTOR-S6K1 signaling cascade are responsible for regulating Atrogin-1 protein content. With the use of our protein content data, other studies should also investigate whether E3 ligase activity of MuRF1 and/or Atrogin-1 is associated with their protein content, and thus a measurement of protein content can be used as a biomarker for E3 ligase activity or vice versa. Additionally, protein degradation contributes half of the equation to determine protein content (i.e., protein synthesis – protein degradation = protein content). Thus, determining degradation mechanisms of Atrogin-1 and MuRF1 protein contents is also important to modulate protein half-life. Thus, understanding of the degradation mechanisms of Atrogin-1 and MuRF1 is required as an important step towards understanding the underlying mechanisms of skeletal muscle atrophy and manipulating their functional E3 ligase activity.

## 5. Conclusions

Based on the findings from the preset study and the existing literature, we propose potential signaling mechanisms that may be involved in the regulation of Atrogin-1 and MuRF1 protein contents in skeletal muscle (**Fig. 6**). The anabolic Akt signaling, which can be activated by Insulin/IGF-1, is a critical upstream signal to modulate MuRF1 and Atrogin-1 at both gene and protein expression levels. However, Atrogin-1, but not MuRF1, protein content is increased when the rapamycin-sensitive and S6K1 dependent signaling pathways are inhibited. Thus, the regulatory mechanisms of protein content are distinct between Atrogin-1 and MuRF1. Our study provides evidence that Atrogin-1 protein content can be regulated by the rapamycin-sensitive mTOR-S6K dependent signaling pathway. Future studies should determine the underlying mechanisms by which the rapamycin-sensitive mTOR-S6K1 signaling regulates Atrogin-1 protein content.

**Figure 6.**
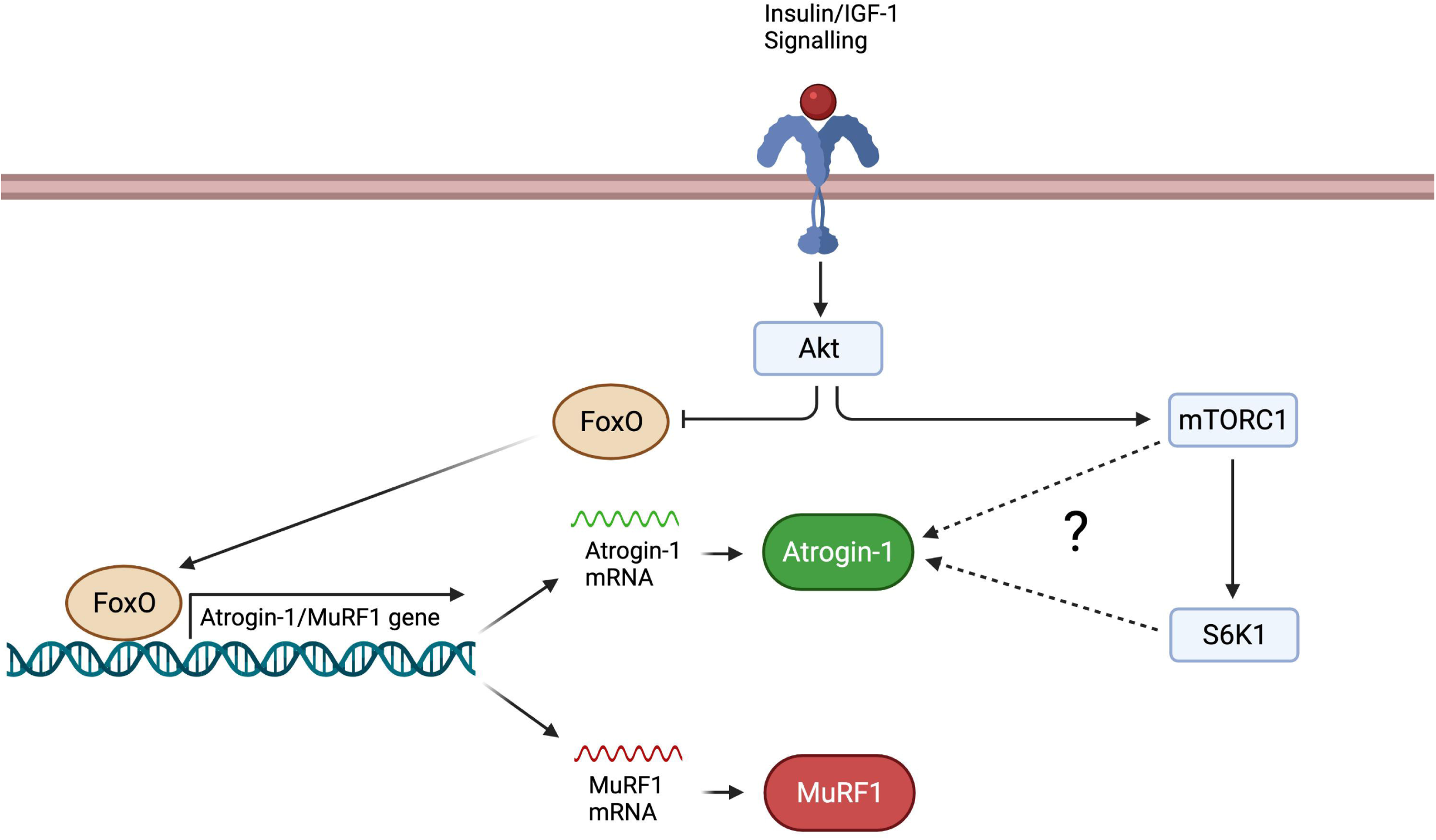
Atrogin-1 and MuRF1 protein contents are differentially regulated in Akt and the rapamycin-sensitive mTOR-S6K1 signaling pathway. Insulin/IGF-1/Akt/FoxO signaling pathway is a predominant mechanism regulating Atrogin-1 and MuRF1 expression at both mRNA transcription and protein levels in skeletal muscle. Upon insulin or IGF-1 stimulation, the binding of their respective receptors triggers a signaling cascade to activate Akt. Akt phosphorylates and inhibits FoxO by preventing their localization to the nuclei, and thus FoxO remains in the cytoplasm. In catabolic conditions, FoxO is less phosphorylated and remains in the nuclei to promote Atrogin-1 and MuRF1 mRNA transcription and thereby increasing their protein content. Inhibition of mTORC1 or S6K1, one of the Akt downstream signaling, can promote Atrogin-1, but not MuRF1, protein content without altering Akt and FoxO phosphorylation. The evidence indicates that Atrogin- 1 and MuRF1 protein content are regulated by at least two different mechanisms. How rapamycin-sensitive mTOR and S6K dependent signaling pathway regulate Atrogin-1 protein content remains undetermined.

## ACKNOWLEDGMENT

Fig. 6 was created with BioRender.com.

## CRediT authorship contribution statement

Yusuke Nishimura: Conceptualization, Investigation, Visualization, Writing - original draft, Writing - review & editing. Ibrahim Musa: Investigation, Writing - review & editing. Peter Dawson: Writing - review & editing. Lars Holm: Writing - review & editing. Yu-Chiang Lai: Conceptualization, Writing - review & editing.

## Abbreviations

MuRF1: Muscle-specific RING finger protein 1
MAFbx: Muscle atrophy F-box protein
PI3K: phosphoinositide 3-kinase
FoxO: forkhead box
mTORC: mechanistic target of rapamycin complex
TSC2: tuberous sclerosis complex 2
S6K: p70 ribosomal S6 kinase
AMPK: adenosine monophosphate activated protein kinase
SDS-PAGE: sodium dodecyl sulfate– polyacrylamide gel electrophoresis
PVDF: polyvinylidene fluoride
TBS-T: Tris-buffered saline Tween-20
ANOVA: analysis of variance
CON: control

